# Geometric purity, kinematic scaling and dynamic optimality in drawing movements beyond ellipses

**DOI:** 10.1101/737460

**Authors:** Adam Matic, Alex Gomez-Marin

## Abstract

Drawing movements have been shown to comply with a power law constraining local curvature and instantaneous speed. In particular, ellipses have been extensively studied, enjoying a 2/3 exponent. While the origin of such non-trivial relationship remains debated, it has been proposed to be an outcome of the least action principle whereby mechanical work is minimized along 2/3 power law trajectories. Here we demonstrate that such claim is flawed. We then study a wider range of curves beyond ellipses that can have 2/3 power law scaling. We show that all such geometries are quasi-pure with the same spectral frequency. We then numerically estimate that their dynamics produce minimum jerk. Finally, using variational calculus and simulations, we discover that equi-affine displacement is invariant across different kinematics, power law or otherwise. In sum, we deepen and clarify the relationship between geometric purity, kinematic scaling and dynamic optimality for trajectories beyond ellipses. It is enticing to realize that we still do not fully understand why we move our pen on a piece of paper the way we do.

**Highlights:**

- Several curves beyond ellipses have power-law **kinematics** with 2/3 exponent.
- The curvature spectrum of each of such **geometries** is quasi-pure at frequency 2.
- Their **dynamics** are shown to comply with minimum of jerk.
- But the 2/3 power law is *not* an outcome of minimizing **mechanical work**.
- Yet, **equi-affine displacement** is invariant upon different kinematics.

**Graphical abstract:** 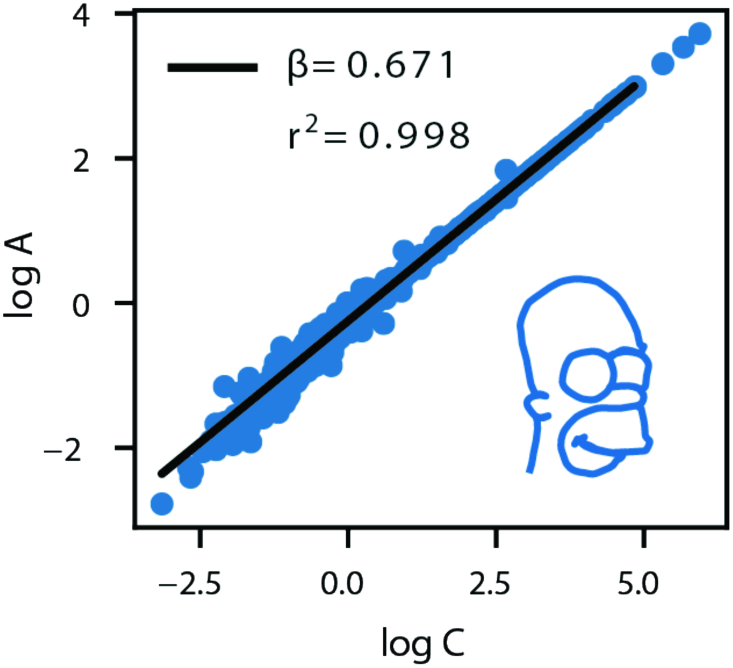

*“We must represent any change, any movement, as absolutely indivisible.” — Henri Bergson*

## 1. INTRODUCTION

In 1609 Kepler published in the book *Astronomia Nova* (**Kepler, 1609**) his celebrated First Law of planetary motion: Mars moves along an elliptical trajectory with the sun at one of its foci. This left behind Ptolomaic and Copernican models; not circles, but ellipses. In the same book we find Kepler’s Second Law, which specifies an invariant (which was later understood as conservation of angular momentum): the area of between the Sun, Mars and any previous point of Mars is constant along the motion of the planet. In sum: equal areas in equal times. This was generalized to all other planets. We move faster when close to the sun (fastest when nearest, at the perihelion), and slower when far away (slowest when furthest, at the aphelion).

Ten years later, Kepler published in *Harmonices Mundi* (**Kepler, 1619**) his Third Law of motion: the semi-major axis A is related to the period P of a planet by means of the following relation: A=k· P^2/3^ (the parameter k is a constant, which can be renormalized by using the Earth’s semi-major axis and number of years as units). It was Kepler’s big achievement to establish such a lawful regularity despite the fact that nobody understood why planets would care to follow it. No one could derive Kepler’s celebrated two-thids power law until Newton’s Law of Universal Gravitation (**Newton, 1687**) was proposed nearly seventy years later. From geometric properties and kinematic laws one would then strive to “climb up” in order to establish dynamic laws that frame the former.

Physics is full of celebrated examples of this sort, where constraints of motion are first discovered and later explained by other more general empirical laws, which in turn are then shown to derive from even more fundamental theoretical principles. Such is a hallmark understanding phenomena, from the motion of planets across the solar system to the movement of Picasso’s brush along a canvas (*in preparation*). However, when it comes to biology, the zeitgeist is mechanistic. The explanatory work seems to be done when a molecule or a circuit is shown to be “necessary and sufficient” for the appearance (or disappearance) of the phenomenon under investigation (**Gomez-Marin, 2017**).

In the midst of the reductionistic zeitgeist obsessed with efficient mechanical causes in the form of counterfactual reasoning within purely interventionist approaches (**Krakauer et al. 2017**), it is conceptually refreshing (and empirically exciting) to realize that relationships like Kepler’s laws can be understood as formal causes. Science is actually the art of interpreting correlations, be it in terms of efficient causation or, in arguably more mature sciences, by actually giving up causation (or, rather, by framing it in) the notion of invariance (**Bailly & Longo, 2011**). Isn’t it ironic that, while the stone falls for symmetry reasons, the insect is thought to fly for neural reasons?

Scaling laws are a particularly relevant sub-class of deep relations, ranging from physics to psychophysics, ecology or language. They all point to unifying principles in complex systems (**West, 2011**). Note that not all power laws are statistical; some relate one degree of freedom to another (like the speed-curvature power law studied here), rather than expressing the functional dependency of a probability distribution.

In curved hand movements, the instantaneous angular speed also scales with local curvature via a power law, whose exponent is 2/3 (**Lacquaniti et al., 1983**). This relationship, simple as it seems, is not a trivial mathematical fact nor is it given by physics (**Zago, Matic et al., 2017**). Cortical computations have been proposed as the controlling mechanism (**Schwartz, 1994**). However, it is still unclear how the neuro-musculo-skeletal system may actually do so. Moreover, the trajectories of insects also comply with the speed-curvature power law (**Zago, Matic et al., 2016**), suggesting that a much simpler explanation —perhaps via simple central pattern generators— may be at work (at least in the humble fruit fly). Nearly forty years later, the origins of the law remain debated.

Most theoretical but also phenomenological studies of the power-law have concentrated on ellipses, also decomposing scribbling into monotonic segments (**Lacquaniti et al., 1983**). On a few occasions shapes other than ellipses have been studies, such as the cloverleaf, lemniscate or limaçon (**Flash et al., 2018**). Invoking optimality as a normative explanation, one can derive the power law by powerful mathematical frameworks. Requiring that the trajectory produces minimum jerk (jerk is the time derivative of acceleration, or equivalently the second derivative or speed, or the third derivative of position) naturally implies such speed-curvature constraints (**Flash & Hogan, 1985**). Also recently, a spectrum of power laws with different exponents has been empirically demonstrated upon drawing a whole range of “pure frequency” curves beyond ellipses, and shown to theoretically derive from minimization of jerk (**Huh & Sejnowski, 2015**).

Notably, it has also been proposed that the 2/3 power-law is an outcome of the least action principle, namely, that imposing mechanical work to be minimal along the trajectory naturally produces the power law with its well-known 2/3 exponent (**Lebedev et al., 2001**). Here we correct such mistaken statement, which allows us to deepen into the relationship between geometrical purity, kinematic scaling and dynamic optimality beyond elliptical trajectories. Planets do not move at constant speed along their (quasi) elliptical trajectories around the Sun. Nor does your finger when tracing an ellipse on a tablet (**Matic & Gomez-Marin, 2019**). And yet, while planets do not follow the speed-curvature power law (**Zago, Matic et al., 2016**), nor do finger movements derive from the physical principle of least action, as we hope to show in what follows.

## 2. MATERIALS AND METHODS

### 2.1. Mathematical calculations

#### Basic notation and equations

Let us use the following notation: A is the angular speed (A=V/R), where V is the instantaneous speed (the module of the velocity vector) and R is the local radius of curvature. Curvature is then defined as C=1/R. The speed-curvature power law then reads: A=k· C^BETA^, where k is a constant and BETA is the power law exponent. By definition, the power law can also be written as V=k· C^BETA-1^.

#### Space-time dilation for arbitrary power-law generation

Since V=ds/dt (where dt is the time differential, and ds the arc-length differential), then one can obtain an explicit relation for how time dilates with space at every infinitesimal increment along the trajectory: ds/dt=k· C^BETA-1^. Since C can be numerically calculated as dtheta/ds, we arrive at the final equation that allows to transform any trajectory into a power law kinematics that respects the original geometry: dt = k^-1^ ds^BETA^ dtheta^1-BETA^.

### 2.2. Numerical simulations

#### Trajectory generation

Trajectories were generated by numerically integrating (with a dt=0.001s) the x and y positions and their derivatives for curves expressed and governed via the following differential equation: d^3^x/dt^3^+x· q(t)=0 for x(t), and also for y(t): d^3^y/dt^3^+y· q(t)=0. Note that the initial conditions can be different but both x and y are governed by the same equation with the same time-dependent coefficient q(t). The four different curves explored in this manuscript were generated by choosing the function q(t) as follows: q(t)=1 corresponds to the ellipse, q(t)=t for the spiral-like ellipse, q(t)=|sin(t)| for “wobbly” curve, and q(t)=|3sin(4t)| for the flower-like curve (see **Figure 1A**).

**Figure 1.**
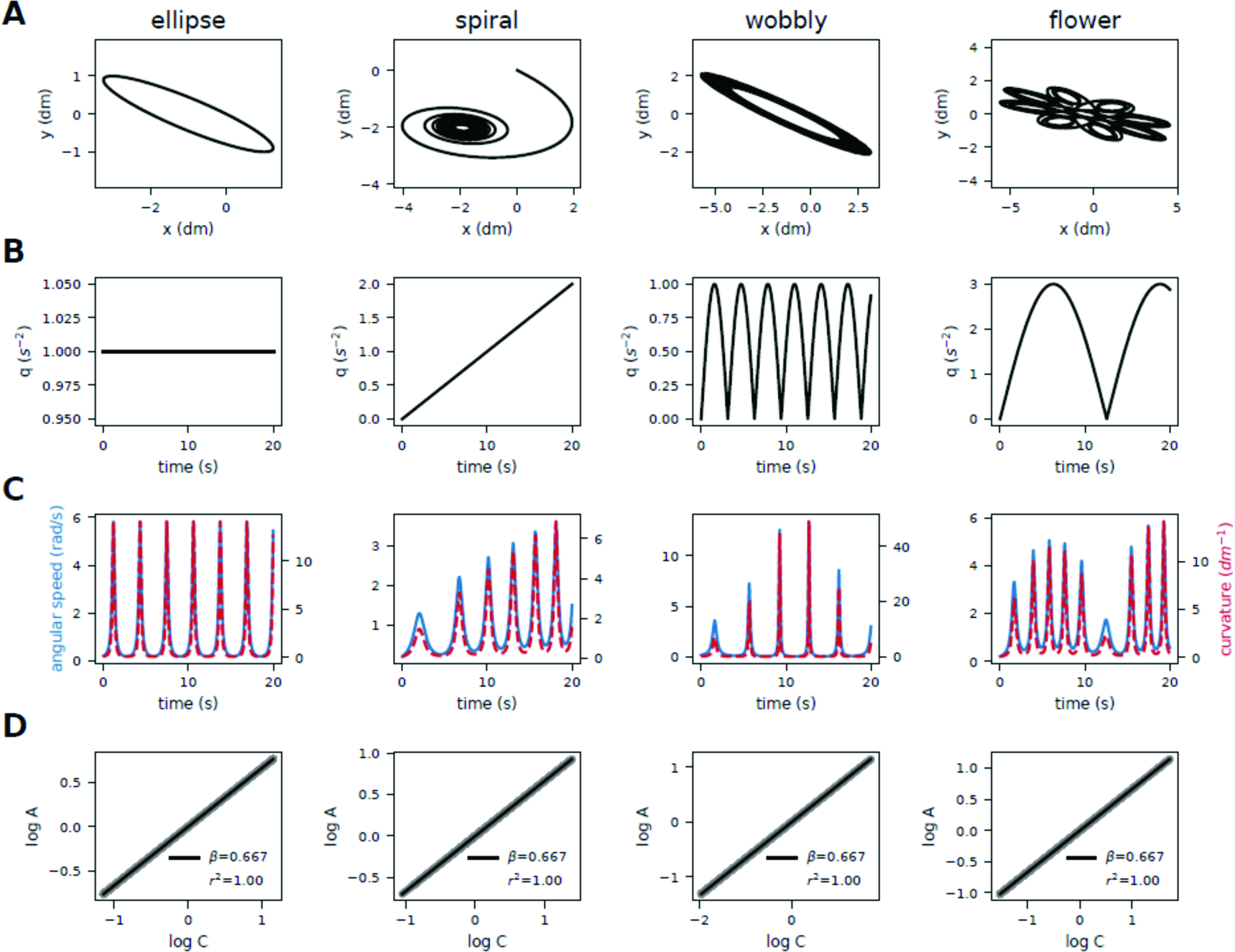
A wide range of geometries beyond ellipses can have two-thirds power law kinematics. (**A**) The four main generated trajectories analyzed in this work: ellipse, elliptic spiral, wobbly ellipse and elliptic flower. (**B**) Time dependence of the function q(t), which generates those trajectories via the third order differential equation d^3^u/dt^3^+u· q(t)=0 satisfied for both x(t) and y(t). (**C**) Time course of instantaneous angular speed A and local curvature C for each curve. (**D**) The numerically estimated log-log plot of angular speed versus curvature reveals, as predicted, an exact power law relationship with exponent 2/3 for each of the curves. Thus an ellipse is not the only geometry that naturally admits kinematic scaling.

#### Curvature spectrum

Curvature frequency spectrum analysis is based on (**Huh and Sejnowski, 2015**), expanded to approximate also the frequency spectrum of non-monotonic angle profiles. We calculate the first derivative of the unwrapped local angle profile, then take its absolute value, and find the anti-derivative. This anti-derivative profile is re-sampled to a uniform step in the local angle coordinate. We take the log of the profile, de-trend it, and apply the Fourier transform.

#### Generating power-law kinematics of any exponent from arbitrary geometries

Selecting an arbitrary power law between angular velocity and curvature is solved by recalculating the time period between each point of the discretized curve, so that the angular velocity fits a desired relationship with curvature (the power-law relation; A=k· C^BETA^), or equivalently, that tangential velocity fits equation (V=k· C^BETA-1^). First, we sample or construct the trajectory using constant step in time (dt). We calculate the arc-length ds_i_, and curvature C_i_ at each point (x_i_, y_i_) of the trajectory. Next we construct a new time-difference vector, where each dt_i_ follows equation dt=(ds/k)· C^1-BETA^. We then construct a time vector T as a cumulative sum of all dt_i._. Next, using a cubic spline, we fit the existing (x, y) points to times T. Then we sample the splined trajectory again with constant dt, obtaining new vector of points (x_i_, y_i_) as a discrete approximation of an arbitrary power law trajectory. Modifying the parameter k then sets the total time of traversing the trajectory without changing the power law relationship.

### 2.3. Behavioral experiments

#### Ellipse trace

Using data from a previous study (**Matic & Gomez-Marin, 2019**), one of the authors traced an ellipse on an android tablet device in a fast and fluid manner. The data was recorded at 85Hz. Raw data was smoothed with a low-pass, 2^nd^ order Butterworth filter, with a cutoff at 8Hz.

#### Homer’s trace

A member of the lab traced a contour of Homer Simpson’s head shown on a Wacom Cintiq interactive graphics monitor, using an electronic pen. The tracing movement was done without lifting the pen from the screen. Several practice traces preceded the trace used in this paper. The data was recorded at 150Hz. Raw data was smoothed with a low-pass, 2^nd^ order Butterworth filter, with a cutoff at 8Hz.

For more details (in fact, virtually all details), please see the **Supplementary material** at the end of the manuscript for the code that generates all analyses and plots.

## 3. RESULTS

### 3.1. A wide range of curves beyond ellipses naturally lead to a 2/3 power law

The speed-curvature power law is the relation A=k· C^BETA^, where A is the instantaneous angular speed (defined as A=V/R) and C is the local curvature (defined as C=1/R), and V is the absolute instantaneous speed of movement and R the local radius of curvature of the trajectory. The term k is a proportionality factor that remains more or less constant empirically (and a precise constant theoretically), and BETA is the power law exponent. This relation is non-trivial since aspects of geometry (like curvature; which concerns only space) and aspects of kinematics (like speed; which concerns time) need not constrain one another in general (like in the motion of a pendulum).

Using the definition of the radius of curvature R as a function of the time derivatives of the trajectory (we are always referring to movement in two dimensions here), it is not difficult to show that, if the power law holds, the term k=D^1/3^, where D=|v_X_ a_Y_ - v_Y_ a_X_|. Obviously, v_i_ and a_i_ are the velocity and acceleration components in both orthogonal directions x and y. The 2/3 power law is often written as A=D^1/3^C^2/3^, with D constant.

Now, if k is constant (namely, if the 2/3 power-law holds), then the term |v_X_ a_Y_ - v_Y_ a_X_|should also be constant. This implies that its time derivative should be zero, and thus one gets: a_X_ a_Y_ + v_X_ j_Y_ - v_Y_ j_X_ - a_Y_ a_X_ = 0 (where “j”, known as jerk, is the time derivative of acceleration; just as “a” is the time derivative of speed). Two terms cancel out, and thus we finally get that any trajectory that complies with the 2/3 power-law must satisfy the following differential equation: j_X_/v_X_=j_Y_/v_Y_. This geometric-kinematic constraint is very interesting because it dictates that both x(t) and y(t) must behave so that the ratio of their third and first time derivatives is equal which, without losing generality, can be expressed as j/v = q(t), where q(t) is any arbitrary temporal function. In other words, one can choose any q(t) at will and, by means of the equation d^3^u/dt^3^+u· q(t)=0 —where u(t) here denotes both x(t) and y(t), although initial conditions can be different—, generate geometric curves whose kinematics follow the 2/3 power law.

Following these mathematical reasoning (**Lebedev et al., 2001**), we generated four different trajectories (**Figure 1A**). Selection of the q(t) function determines the shape of the trajectory: for the ellipse, it is constant, q=1; for the elliptic spiral q=t; the wobbly ellipse, q =|sin t|; and for the elliptic flower we chose q=|3 sin t/4| (**Figure 1B**). Not only are curvature and angular speed of these trajectories strongly correlated (**Figure 1C**), they in fact follow the 2/3 speed-curvature power law exactly (**Figure 1D**).

**Supplementary Figure 1.**
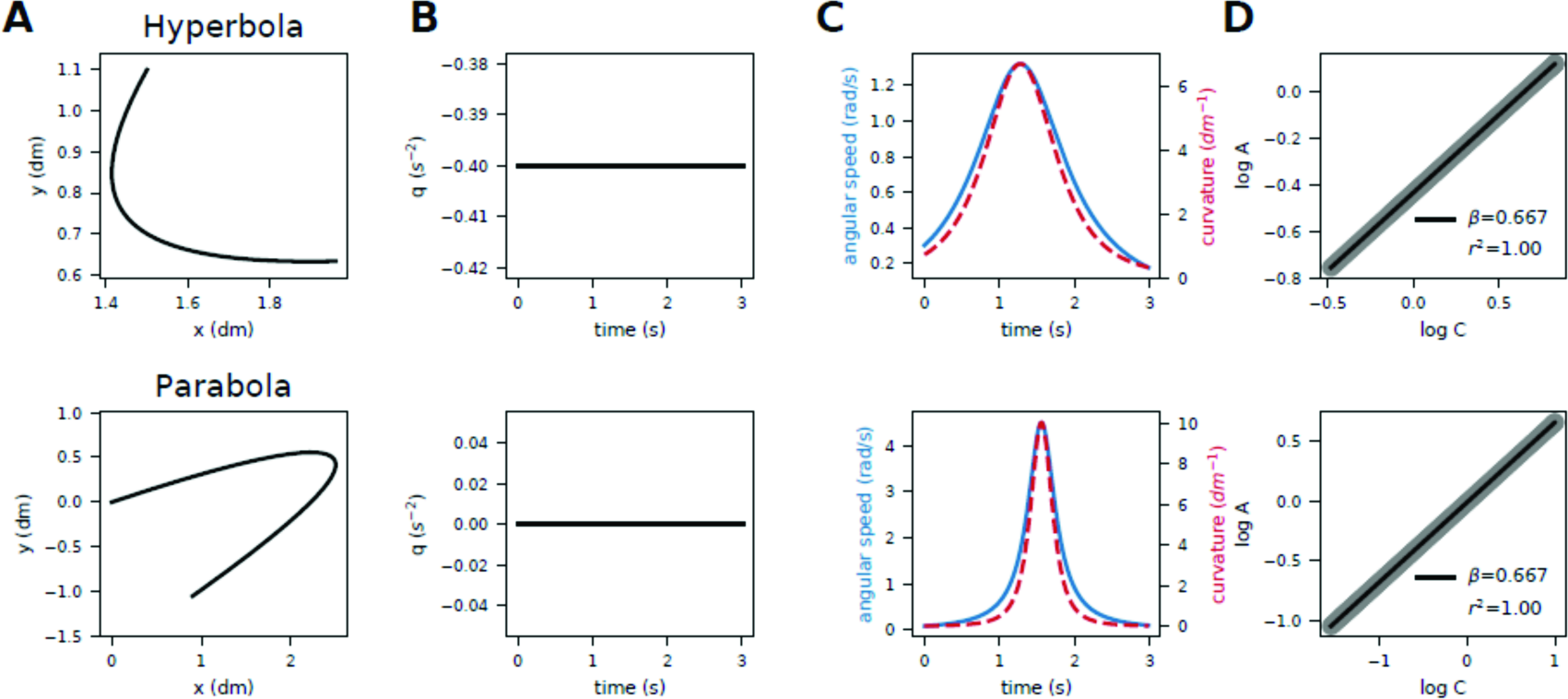
Apart from the ellipse, the hyperbola and the parabola are generated with constant q, and also lead to 2/3 power-law kinematics. (**A**) Both curves. (**B**) Function q(t) that generates the curves. (**C**) Speed and curvature in time. (**D**) Log-log plot of speed and curvature, numerically demonstrating an exact power law constraint.

Lebedev and colleagues explicitly listed the ellipse, hyperbola and parabola as the trajectories resulting from constant q(t), noting the relationship between constant q and the resulting geometry (q=0 for parabola, q < 0 for ellipse, and q > 0 for hyperbola). In **Supplementary Figure 1** we analyzed those three curves in the same way as four mentioned curves in **Figure 1**. Curvature and angular speed are visibly constrained, following a power law with the exponent of exactly 2/3.

To the best of our knowledge, nobody has analyzed the family of curves that are generated with a non-constant q(t), some of whose exemplars we show in **Figure 1**. In what follows, we will concentrate on such four curves to gain further insights into geometric purity, kinematic scaling, and dynamic optimality. We will also correct an important physics error in (**Lebedev et al., 2001**).

### 3.2. Two-thirds power law trajectories have quasi-pure geometrical spectra

Let us now concentrate on the geometry of the curves presented in the previous section. It has been recently shown that the speed curvature power laws (of different exponents, not just 2/3) are achieved for so-called “pure frequently curves” (Huh and Sejnowski, 2015). Actually, (as we will see in the last section of the Results) trajectories with “mixed curvature frequencies” cannot comply with the kinematic scaling of the power law, unless they give up dynamic optimality. So, how does one estimate the “geometric purity” of a curve?

Parametrization of local curvature can be done in many ways. To estimate power laws one usually parametrizes curvature in time, namely, C(t), so that it can be compared, moment to moment, with speed V(t), which is naturally defined as a function of time. Time parametrization of log curvature is convenient in the regression analysis with log angular velocity, also parametrized in time (as in **Figure 1D**). In **Figure 2B** we show curvature parametrized in time for the four study-case trajectories shown in **Figure 2A**.

**Figure 2.**
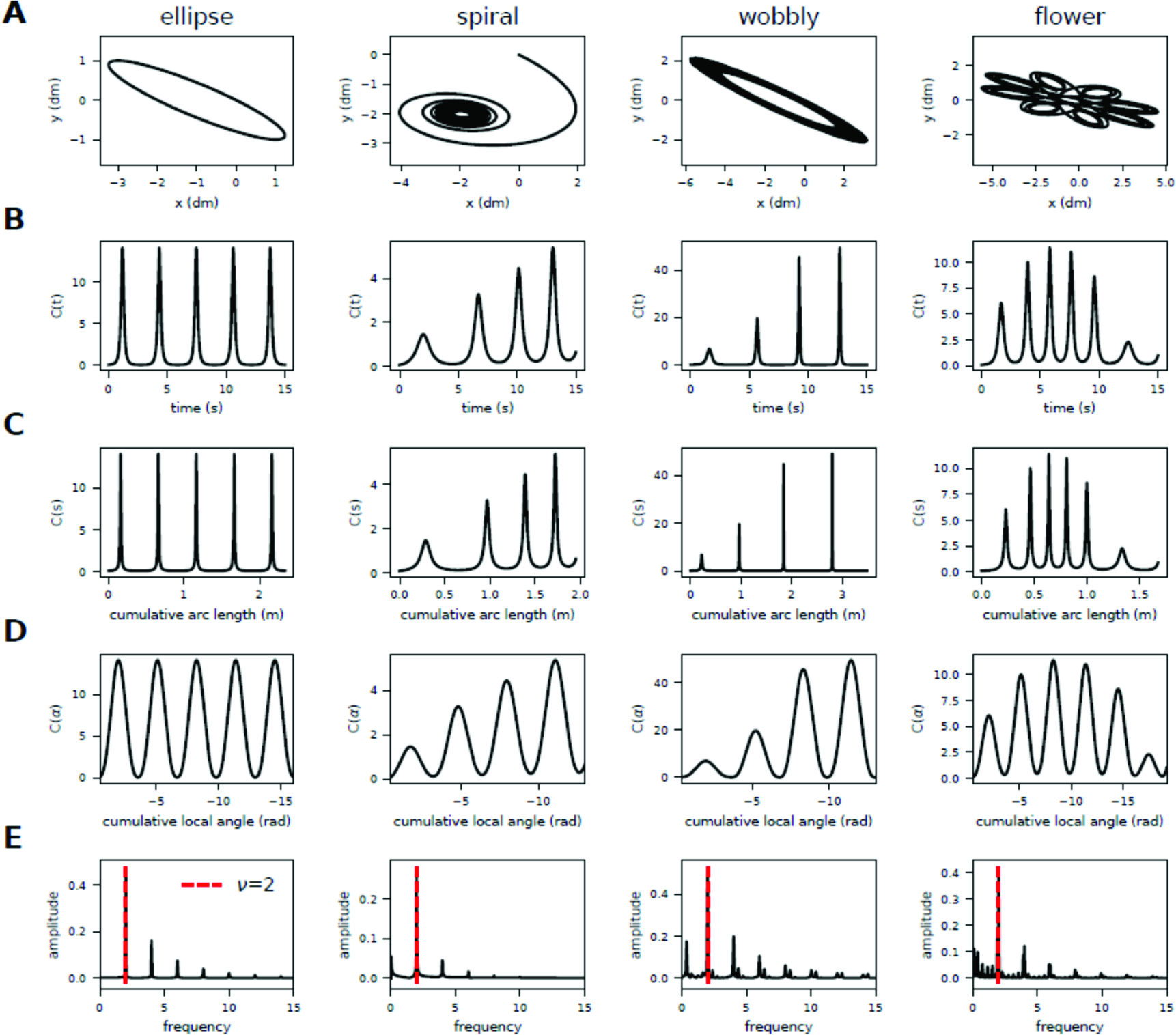
Very different geometries can have the same quasi-pure curvature spectrum. (**A**) The four studied curves in Euclidean space. (**B**) Their local curvature parametrized in time “t”. (**C**) Curvature parametrized in arc length space “s”. (**D**) Curvature reparametrized in local angle “alpha”. (**E**) Geometric spectrum of the curves —Fourier transform of the logarithm of C(alpha)—, showing that all curves are quasi-pure with a dominant peak at frequency 2.

However, cumulative arc length (s) is the natural parametrization for curvature, since curvature is by construction a purely geometrical quantity, and so the time parametrization natural in kinematic quantities (such as speed) injects a temporal bias that geometry should be indifferent to. In **Figure 2C** we re-parametrize curvature now in terms of arc length, C(s). Note the subtle change in the functions with respect to the time parametrizations in **Figure 2B**.

There is a third way to parametrize curvature: rather than time or length, one can use angle. Based on (**Huh, 2015**) we can parametrize curvature in the local angle coordinate, as shown in **Figure 2D**. This representation has many advantages in understanding essential properties of the curves, as well as revealing the connection between geometry and kinematics in power law constraints (**Huh and Sejnowski, 2015**).

In particular, once any curve is parametrized in the angle, one can detect a shared feature in the four curves studied here: note how the profiles of **Figure 2D** have naturally rescaled with respect to those in **Figure 2C** and **Figure 2B**, so that now, every 2π, curvature undergoes exactly two complete oscillations. If these were temporal functions, a Fourier transform would immediately reveal a dominant frequency there.

Following (**Huh and Sejnowski, 2015**) we apply the Fourier transform to the log of the curvature profile, once parametrized in the local angle coordinate (**Figure 2E**). The resulting amplitude profile shows curvature frequency spectrum in angle space. The frequency of a curve is the number of curvature oscillations per unit of local angle (full oscillation is 2π radians), and the local angle is defined as the angular direction of the velocity vector. Despite their very different appearance in X-Y space (**Figure 2A**), all four curves share a main peak at exactly ν=2 (which corresponds to Huh’s pure ellipse; see below) as well as some ripples.

The quasi-pure spectrum of these geometries, and specially that of the ellipse shown in on the left side of **Figure 2A**, makes one wonder why they are not exactly pure (namely, with a single peak at ν=2, without any ripples). To better understand this, we went back to Huh’s pure frequency curve with ν=2 (**Huh, 2015**), which is visually very similar to the classical ellipse, (x/a)^2^+ (y/b)^2^=1. Both curves are shown in **Supplementary Figure 2A**, together with ellipses empirically traced on a tablet.

Huh’s ellipse (on the left) has a single strong peak at ν=2 by design, and no peaks at other frequencies, meaning that its log-curvature profile in angle space is a pure sinusoid. The classic ellipse, constructed with two orthogonal sine waves with 90° phase difference, has a few harmonics at frequencies multiples of ν=2 (4, 6, 8, etc), but it is still quasi-pure. The empirically recorded ellipse trace, similarly, shows some harmonics and also peaks at other frequencies (**Supplementary Figure 2B**). It is also decently pure. In sum, this precise geometrical analysis of the spectrum of curvature is both informative as to whether we shall expect a power law and of what exponent, but also a necessary condition to know that we are dealing with a pure frequency curve in the first place, which is very important when trying to determine whether the speed-curvature power law holds empirically.

Finally, to gain even further insight into what these spectra are reflecting, we morphed an ellipse into a circumference by reducing the eccentricity of the former (**Supplementary Figure 2C**). The amplitude of the peak at ν=2 is progressively reduced, as well as all the other harmonic frequencies, until the circumference does not show peaks at any frequency (as it should, since its curvature is constant).

We can proceed now with kinematic and dynamic considerations on these curves.

**Supplementary Figure 2.**
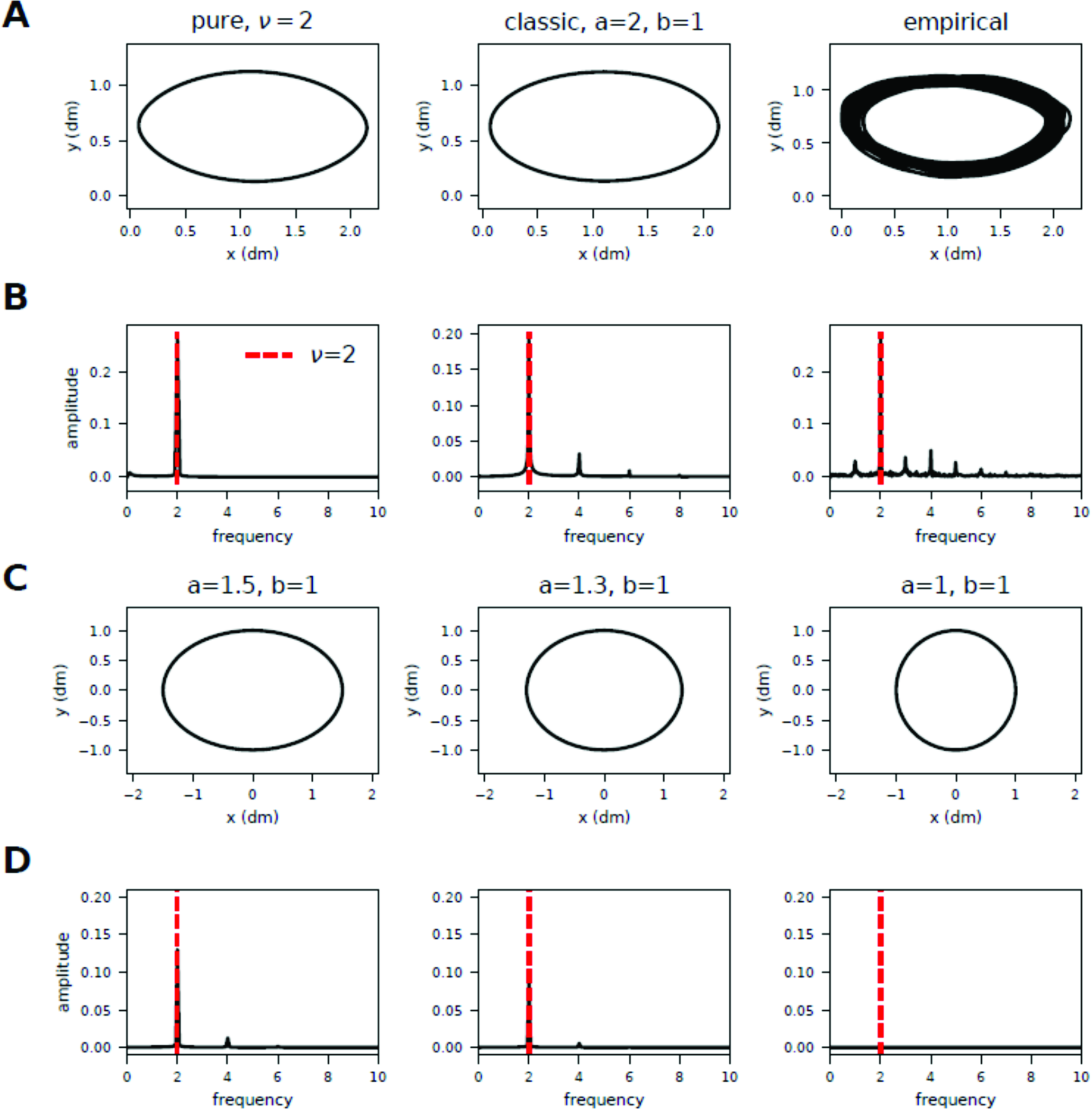
Further exploration of basic properties of curvature spectra for ellipses. (**A**) Three ellipses: a pure frequency curve with ν=2, or “Huh’s ellipse” (left), the “classic” ellipse constructed with two orthogonal sine waves, ninety degrees out of phase (middle), and empirically traced ellipse (right). (**B**) Curvature spectra with the pure peak at ν=2 shown in red. (**C**) Classic ellipses with decreasing eccentricity turning into a circumference. (**D**) Corresponding geometry spectra, showing the peak at ν=2, with the amplitude of the peak decreasing with eccentricity.

### 3.3. The power law does not imply that mechanical work is constant nor minimal

For 2/3 power law trajectories, we have seen that D is constant. It turns out that D is actually the magnitude of the *cross* product between the velocity and acceleration vectors. And so, for the trajectories displayed in (**Figure 3A**) that dot product should be constant too (**Figure 3B**). Such magnitude can be represented as the surface of the parallelogram closed by such vectors (**Figure 3D**). So far, so good.

**Figure 3.**
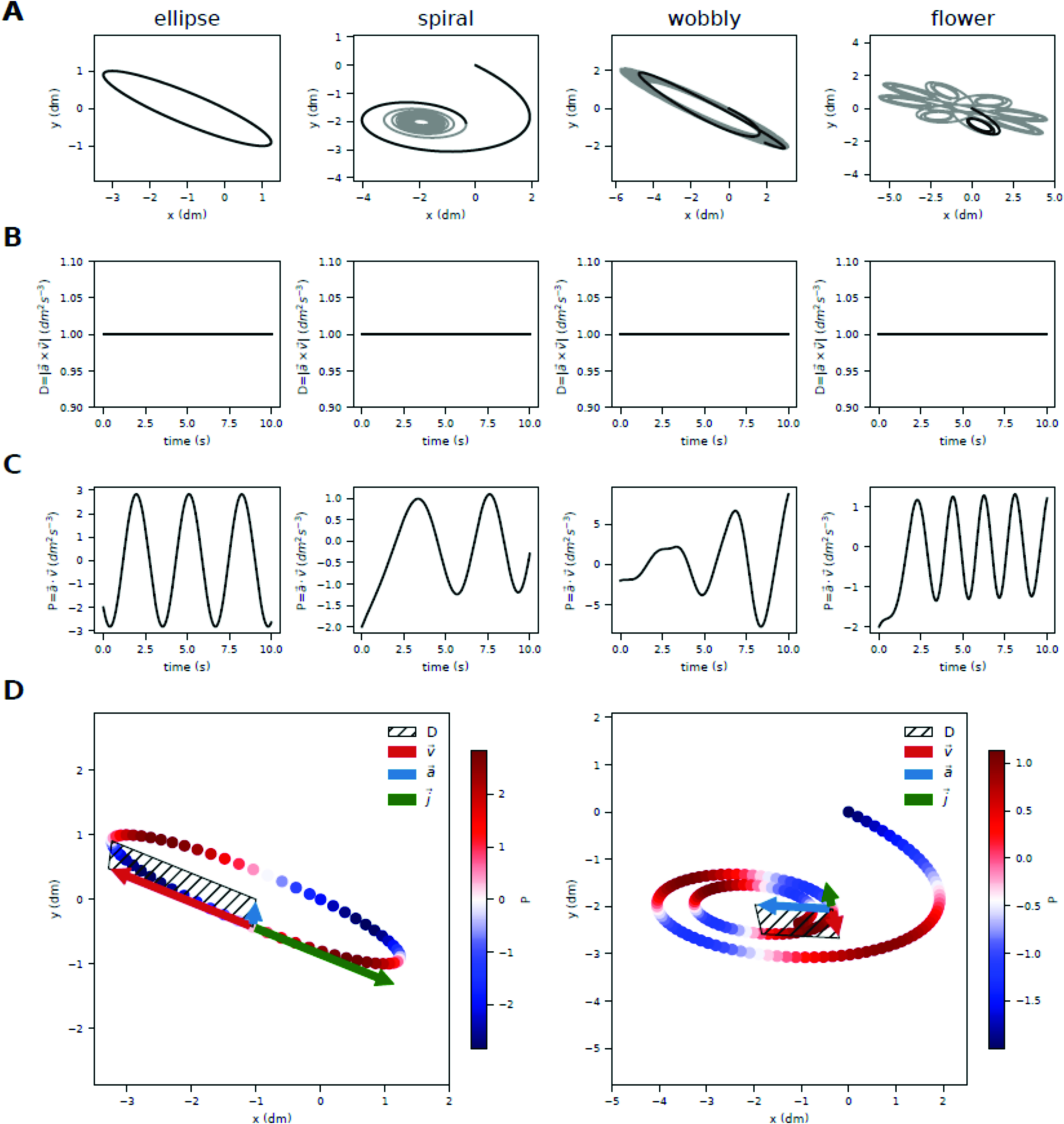
For 2/3 power law trajectories, the term D is constant but mechanical work is not. (**A**) Analyzed segments of the curves marked in solid black. (**B**) Magnitude of the cross product of the velocity and acceleration vectors (which is mathematically equal to D) as a function of time along the trajectory. It is constant for all curves. (**C)** Dot product of the velocity and acceleration vectors (which is proportional to mechanical work) over time. It is not constant for any curve. (**D**) Velocity, acceleration and jerk vectors for two of the curves at a given time instant. Graphically, the term D is the area of the parallelogram formed by velocity and acceleration, and it has the property of remaining constant along the trajectory. The jerk vector also has the property of being anti-parallel to the velocity vector. Mechanical power (depicted in color) is not constant along the trajectory. It is actually zero at the extremes of the ellipse (since velocity and acceleration vectors are orthogonal there) but non-zero at other times. See also animations of panel (D) as **Supplementary Material**.

Remember that one can rewrite the 2/3 power law (A=kC^2/3^) as A=D^1/3^C^2/3^, and then simply as V= D^1/3^R^1/3^, so that V^3^/R=D. With similar mathematical manipulations (**Lebedev et al., 2001**) arrive at this last same equation and, rewriting D=V(V^2^/R) realize that the term in parenthesis is the magnitude of centripetal acceleration (A_n_), and so D=V· A_n_. The fatal error comes in their equation (5), when they say that “[t]his product is known in physics as mechanical power”, which they call P. The essential mistake that invalidates the main claim of their paper is that D=P.

If that was the case, then a 2/3 power law would constraint movement along the trajectory to have constant mechanical power (because we have seen that D is constant). As we will unpack further below, the authors are naturally thrilled to discover that, mathematically, the time integral of D is minimal when D happens to be constant. In other words, the “optimal” way to move is to do so that D is constant, aka, the 2/3 power law. They are thrilled (as we would) because, if the physics were true, the mathematics would prove that “drawing movements [which fulfill the 2/3 speed curvature power law] are “an outcome of the Principle of Least Action” (which is precisely the title of their paper). But if D is not the mechanical power, then the claim evaporates.

Why isn’t D=P, then?

Lebedev and colleagues equate mechanical power with D, namely, the authors take the product of centripetal acceleration with the speed to be proportional to physical force that would push a particle moving along such 2/3 power law trajectories.

Mechanical power is the amount of mechanical work per unit of time. Mechanical work is the amount of energy transferred by a force. It is calculated as the integral of the force vector along the trajectory vector. Force is proportional to acceleration, and the trajectory vector can be rewritten as velocity times dt. Thus, in practice, mechanical work is proportional to the product of velocity and acceleration. But (and here comes the subtle mistake), it is the *dot* product (also called *scalar* product) of the vectors, rather than the simple product of their magnitudes. Put plainly, the dot product of two orthogonal vectors is zero, no matter how large they are; while the product of their magnitudes is large. In sum, mechanical work is calculated via the *scalar* product —rather than the *cross* product (which gives us D)— of velocity and acceleration. And thus, as shown in **Figure 3C**, work is far from constant along the trajectory, as opposed to D (**Figure 3B**).

Animated traversals of the spiral and elliptical trajectories are available in the *github* repository as video files (see **Supplementary Material**). They show the constancy of the magnitude of the cross product D during the whole trajectory, and the changes in the position, acceleration, velocity and jerk vectors over time. See **Figure 3D** for snapshots.

In sum, and contrary to what was claimed in (**Lebedev at al., 2001**): the 2/3 power law is *not* an outcome of the physical principle of least action.

### 3.4. Trajectories with constant D minimize the time integral of D

The mathematical derivation that, by means of a variational analysis, shows that the time integral of D is minimal when D is constant (**Lebedev et al., 2001**) is still valid and somewhat insightful. Agnostic about the existence of a meaningful physical or mathematical interpretation of the term D, next we sought to numerically demonstrate that constant-D prescribes the most “economical” way to move amongst the infinitely many ways to do so. To our knowledge, such minimization hasn’t been done numerically.

Because we seek a numerical demonstration that trajectories complying with the 2/3 power law constrain their geometry (curvature) and kinematics (speed) so as to minimize the integral of D, we can only aspire to show local, rather than global, minima. To that end, we invented a way to systematically generate a range of different kinematics that would transverse the exact same geometry in the exact same total duration (see below).

We take a segment (of a trajectory that complies with the 2/3 speed curvature power law) with starting points A and B (**Figure 4A**), and whose total time duration is T (vertical black line in the plots of **Figure 4B**). We then maintain the geometry but rescale the kinematics so that the same segment is traversed in the same amount of time but now with a kinematics that would still yield a power law but with an exponent different than 2/3 (say, with hypo-natural exponent 1/3, and hyper-natural exponent equal to 1). We then numerically calculate D as we integrate it in time all the way to t=T (**Figure 4B**) for such three different (power law) kinematics. Exponent 2/3 always yields the minimum value at the end of the segment. In **Figure 4C** we can calculate I(t=T) for a whole range of values of β on the given path. Compared to trajectories with any other values of β, trajectories with β=2/3 indeed have the minimal time integral of D.

**Figure 4.**
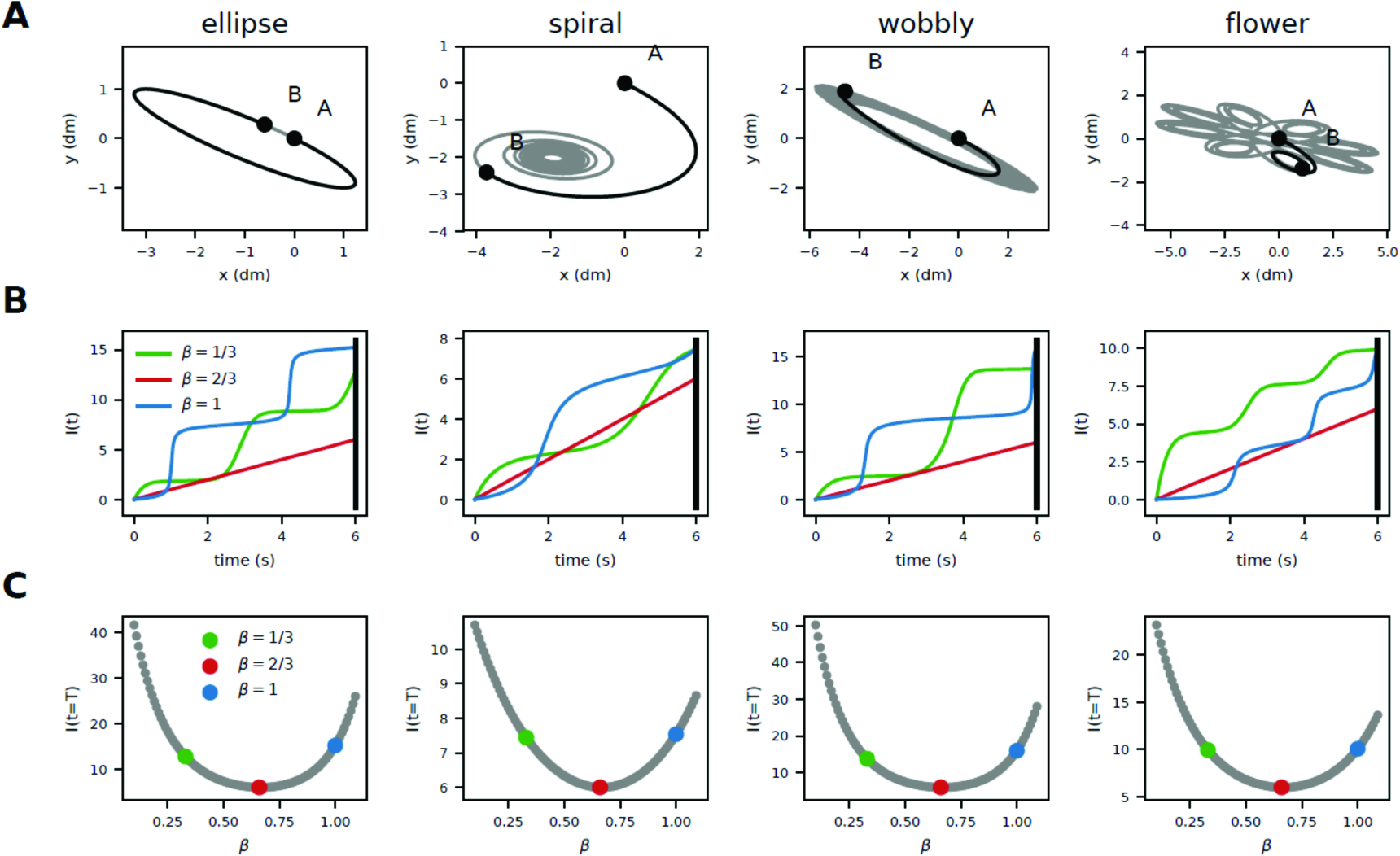
Numerical evidence that the time integral of D is (locally) minimal when D is constant. (**A**) Start and end points of trajectory segments (of 6 seconds of duration) for each of the four main curves. (**B**) The time integral of D as time elapses from the beginning to the end of the segment. Respecting the same geometry and total duration of the movement, different kinematics were explored (generated by power laws with different exponents; depicted in green, red blue). (**C**) The integral of D, when numerically calculated for all exponents between 0 and 1 turns out to be minimal for β=2/3 (red dot), for all curves.

**Supplementary Figure 4.**
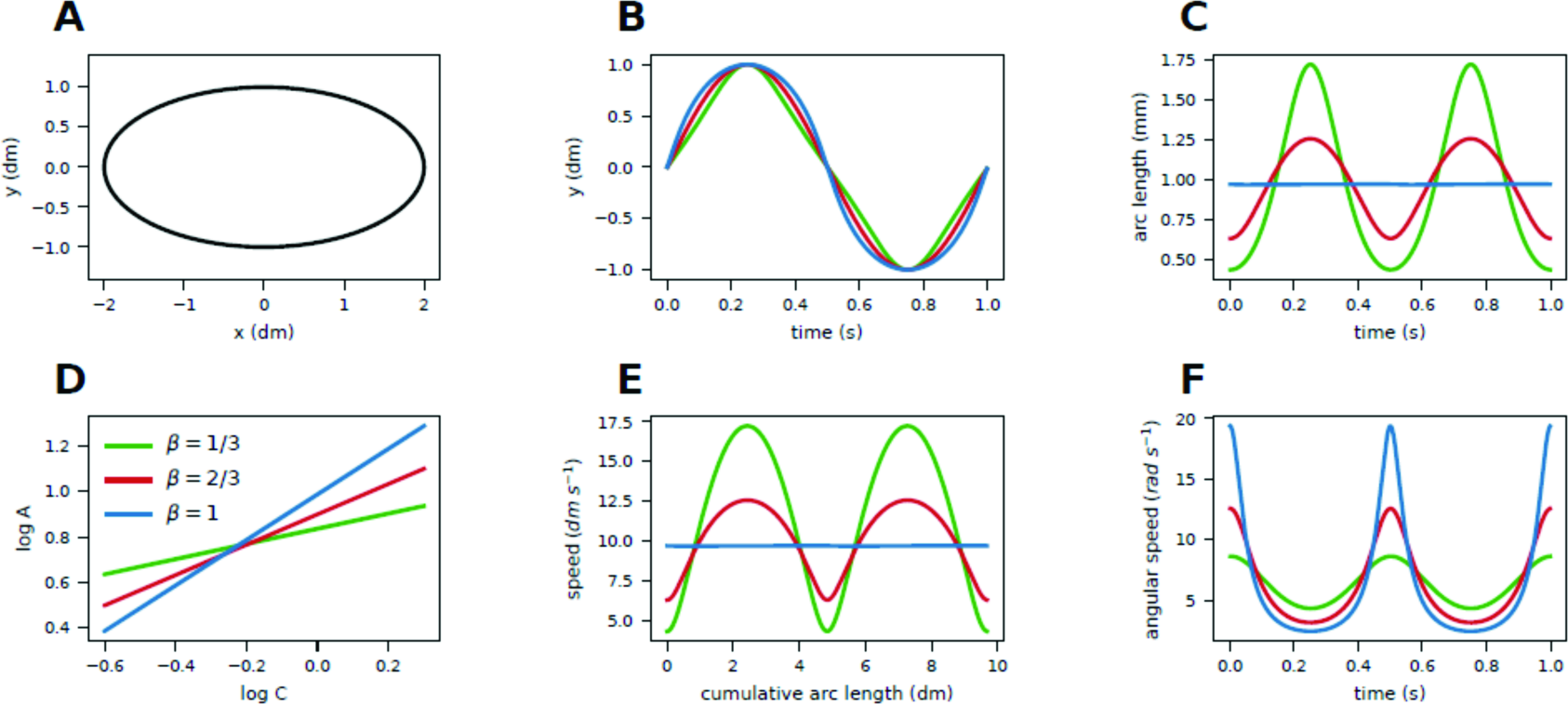
Creating speed-curvature power law trajectories of arbitrary exponents. Given a fixed geometry, a trajectory of any power law exponent can be generated (**A**) A ‘classic’ ellipse geometry (**B**) Y positions in time for trajectories with different exponents. (**C**) The corresponding arc-length rescaling of the original trajectory, and (D) the log-log plot shows the generated power laws with different exponents. (E) Speed over cumulative arc length (path traveled), rescaling for different exponents. (F) Angular speed over time.

In case it is not already clear by now, let us emphasize that a given geometry can in principle be traversed with any kinematics. Let us now have a brief interlude to explain and illustrate how to kinematically re-scale a given geometry with any kinematics to a power-law kinematics with our exponent of choice. For each β in the required range, we start with a generated path as an ordered list of points. Given the path, the β and k, we calculate the time periods between each point of the path, so that they satisfy the formula dt = (ds/k)· C^1-BETA^, derived as explained in the Methods. The resulting trajectory does not necessarily have the desired average speed. The whole trajectory is then re-calculated with the same points and β, but with a different k parameter until the average speed is within tolerance from desired average speed.

We illustrate the effects of such rescaling algorithm for the classical example of an ellipse (**Supplementary Figure 4A**). Generating an elliptical trajectory with orthogonal sine waves yields a β=2/3 power law (**Supplementary Figure 4D**, red line). We can rescale this trajectory into β=1 (blue line) and β=1/3 (green line) power laws. The Y coordinate over time (**Supplementary Figure 4B**) of the β=2/3 trajectory is shown in red, and is a pure sinusoid. A trajectory with exponent β=1 is more ‘round’ in the Y coordinate, and a trajectory with β=1/3 is more ‘triangular’. The arc-length for the β=2/3 trajectory changes over time: an object moving on such trajectory slows down in more curved parts, and speeds up in straighter parts of the path (**Supplementary Figure 4C**). Because human participants produce speed profiles similar to these, the β=2/3 trajectory is called ‘natural’. In comparison, a trajectory with β=1 is called ‘hyper-natural’ and has a constant arc length, because it has constant tangential speed. Trajectories with β=1/3 are called ‘hypo-natural’, as they slow down more and speed up more than β=2/3 ‘natural’ trajectories. Similar relationships are visible in the speed over cumulative arc length plot (**Supplementary Figure 4E**), illustrating the transformations made by the rescaling algorithm. When shortening the time period of crossing the same distance, we get higher speed, as illustrated by the peaks of the β=1/3 (green) plot. For longer times, speed goes down, as in the valleys of the β=1/3 plot. Angular speed over time (**Supplementary Figure 4F**) shows some inverted relationships. Here, the hyper-natural trajectory has highest peaks and lowest valleys of the three trajectories.

### 3.5. Equi-affine displacement is invariant under different kinematics

We have seen how the time integral of the term D is minimal when D is constant. However, we have also seen that D does not correspond to mechanical power, and so the minimization of D does not imply that the power law is the outcome of the least action principle of physics. Is there any other quantity whose integral, when minimized, lends itself to a meaningful interpretation?

The cube root of D has been identified as the so-called equi-affine speed (**Pollick and Sapiro 1997; Flash and Hadzel, 2007**): V_EA_=D^1/3^. Of course V_EA_ is constant when D is constant. But note that the fact that the integral of D is minimal when D is constant does not mean that the integral of V_EA_ is minimal when V_EA_ is constant. What happens when we minimize the integral of V_EA_?

We can answer such question mathematically by means of variational calculus. When deriving Euler-Lagrange equation that results in minimizing the equi-affine speed as the Lagrangian, we found that the terms in such equation cancel out completely. Aren’t there any particular solutions that make the functional an extremum?

We then answered such question numerically. We followed in **Figure 5** the same procedure as in **Figure 4**. We took our four main curves and chose a segment of duration T (**Figure 5A**) and numerically estimated the time integral of V_EA_ upon movement along the same geometry with three different kinematics (**Figure 5B**), this is, power laws with different exponents. To our surprise, and as opposed to the integral of D in **Figure 4**, the integral of V_EA_ yields the same value at the end of the segment (t=T) regardless of the kinematics. There seems to be no minimal. Is it thus an invariant?

**Figure 5.**
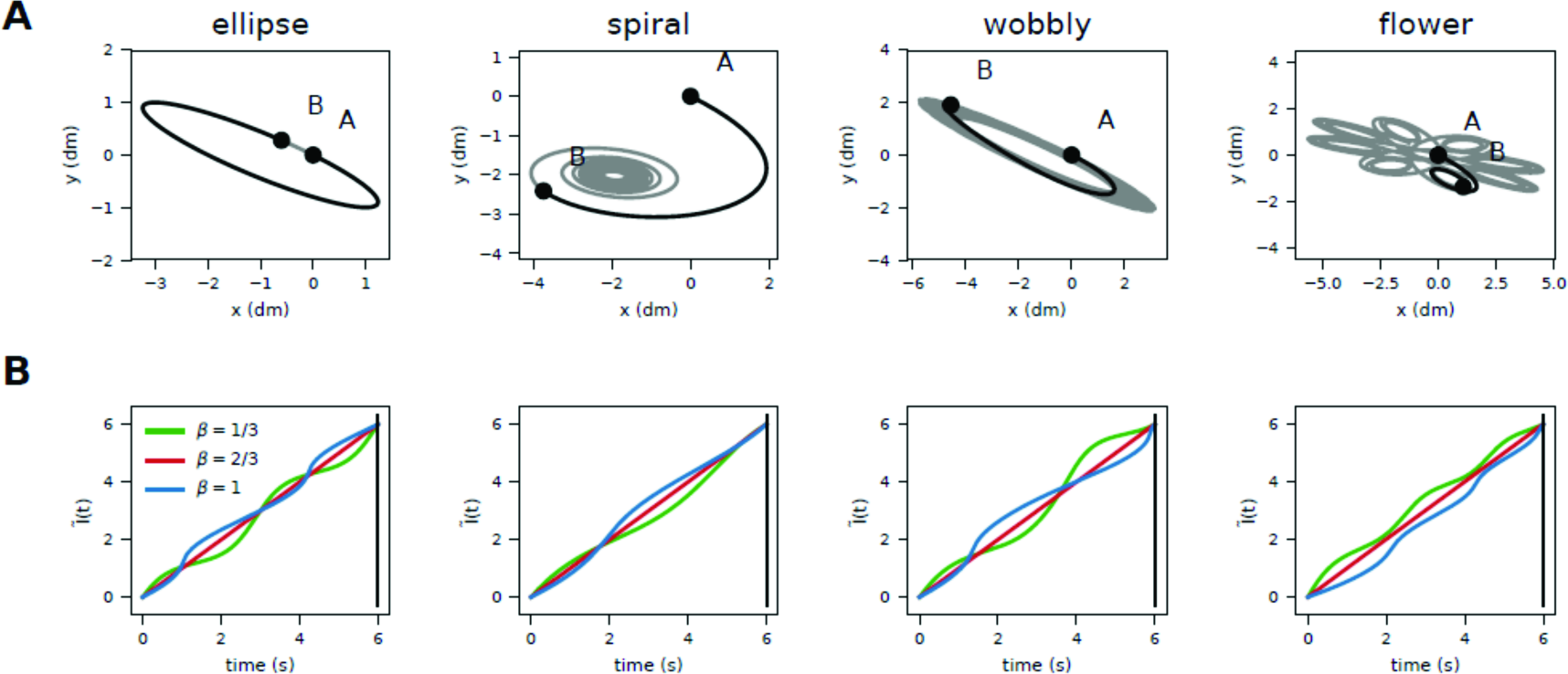
The time integral of equi-affine speed is invariant for different kinematics with fixed geometry. **(A)** Trajectory segments of duration T (in black) with their start and end points. (**B**) Time integral of equi-affine speed for various kinematics (power laws of different exponents). Unexpectedly, all kinematics lead to the same value of the integral at t=T, for all curve segments.

Note that, generally, the time integral of speed along a path is precisely its total displacement. In fact, the integral of affine velocity is known as the equi-affine arc-length or the special affine arc-length (**Izumiya and Sano, 1998**). Our analytical and numerical results thus indicate that affine arc-length is invariant under different power law kinematics.

Next we asked whether such invariance remains when the kinematics does not follow a power law (**Supplementary Figure 5A**) and/or when the geometry between A and B is different (**Supplementary Figure 5B**).

**Supplementary Figure 5.**
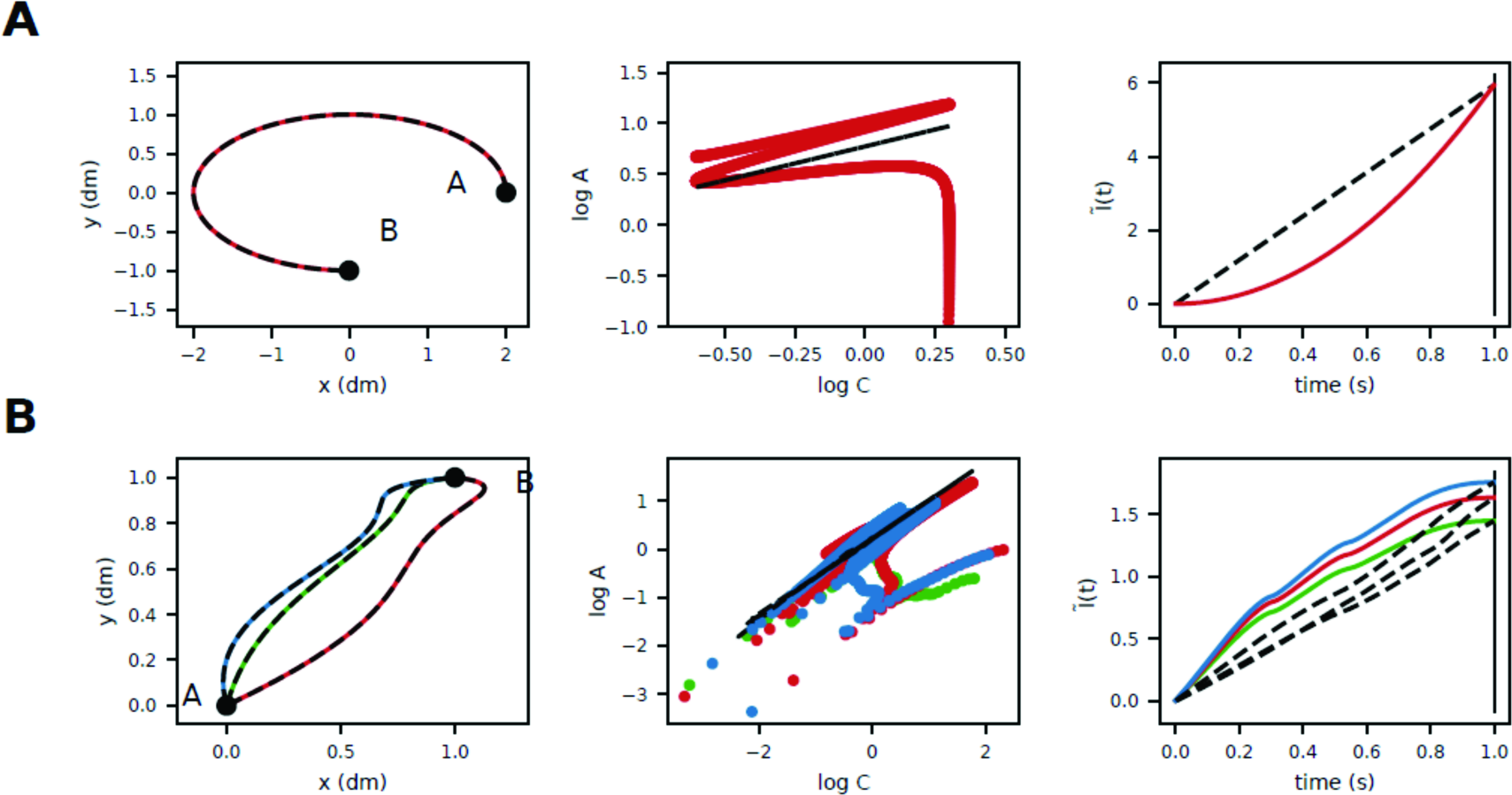
Equi-affine displacement is invariant upon different kinematics traversing the same geometry, but not for different geometries with the same start/end positions. **(A) Left:** Segment of an elliptical path between points A and B of duration T. **Middle:** Its corresponding log-log plot of speed and curvature (middle). Black color indicates that the trajectory follows a power law (PL), while the red one does not (both by construction). **Right:** The time integral of equi-affine speed from t=0 to t=T yields the same value for both PL and non-PL kinematics, given the same geometry. **(B) Left:** Three pseudo-random paths (different geometry) and different kinematics, all going from point A to B in the same time interval T. **Middle:** Dotted lines have power law kinematics, and colored lines non-power law kinematics. The integral of equi-affine speed is the same for power law kinematics and non-power law kinematics.

In a similar analysis to **Figure 5**, we show that the affine arc-length is the same for power law and non-power law kinematics. An elliptical trajectory segment (Figure 5.1 A) is traversed with power law kinematics (with exponent β=2/3) (in black) and non-power law kinematics (with ellipse’s sine angle theta increasing with time squared) (in red). The integral of equi-affine speed is the same for both (**Supplementary Figure 5A**).

Let us note an interesting pathological case: in movement from A to B in a straight line at constant speed, there is no acceleration vector, and so V_EA_ is zero and so is its integral.

To explore the effect of different ways to get from one point to another in space (geometry), not just in time (kinematics), we also tested three pseudo-random paths from points A to B. Using the procedure described in (**Supplementary Figure 4**), we imposed power law kinematics (black lines), while colored lines had non-power law kinematics, as shown in the middle plot. The integral of equi-affine speed is the same for both kinematics, but not across different geometries (**Supplementary Figure 5B**).

Equi-affine speed is not invariant under arbitrary transformations. It has been shown that equi-affine length is invariant under affine transformations using the signed volume of the parallelepiped created by vectors of first, second and third derivative with respect to time of the curve r, raised to the power 1/6 (**Pollick et al., 2009**). Equi-affine speed has also been shown to be piecewise constant along movement segments and so, rather than Euclidian, it becomes a natural geometric description of hand trajectories (**Flash and Handzel, 2007; Polyakov et al., 2009; Bennequin et al., 2009; Meirovitch et al., 2016**). However, we have not been able to find an explicit claim that the time integral of equi-effine speed is a kinematic invariant, as our findings suggest.

### 3.6. Pure curves with two-third power law scaling minimize jerk

Having found a way to numerically estimate whether certain functionals (such as D and V_EA_ respectively in **Figure 4** and **Figure 5**), are (locally) minimal for a fixed geometry upon different kinematics, we now apply the method to confirm (**Huh and Sejnowski, 2015**) mathematical derivations: minimum of total jerk is achieved for pure frequency curves when their kinematics follow a speed-curvature power law (where the exponent value β depends on the frequency ν of the curve).

It is well known now that ellipses (which we have shown to have ν near to 2), when traversed with a power law kinematics of β=2/3 (which is how they are traced by humans), have minimum jerk (**Wann et at, 1988; Viviani and Flash 1995; Huh and Sejnowski, 2015**). But, our knowledge, nobody has estimated this numerically. Nor has this claim been shown for the large family of curves that, despite not being an ellipse, have ν=2 (like those in **Figure 6A**).

**Figure 6.**
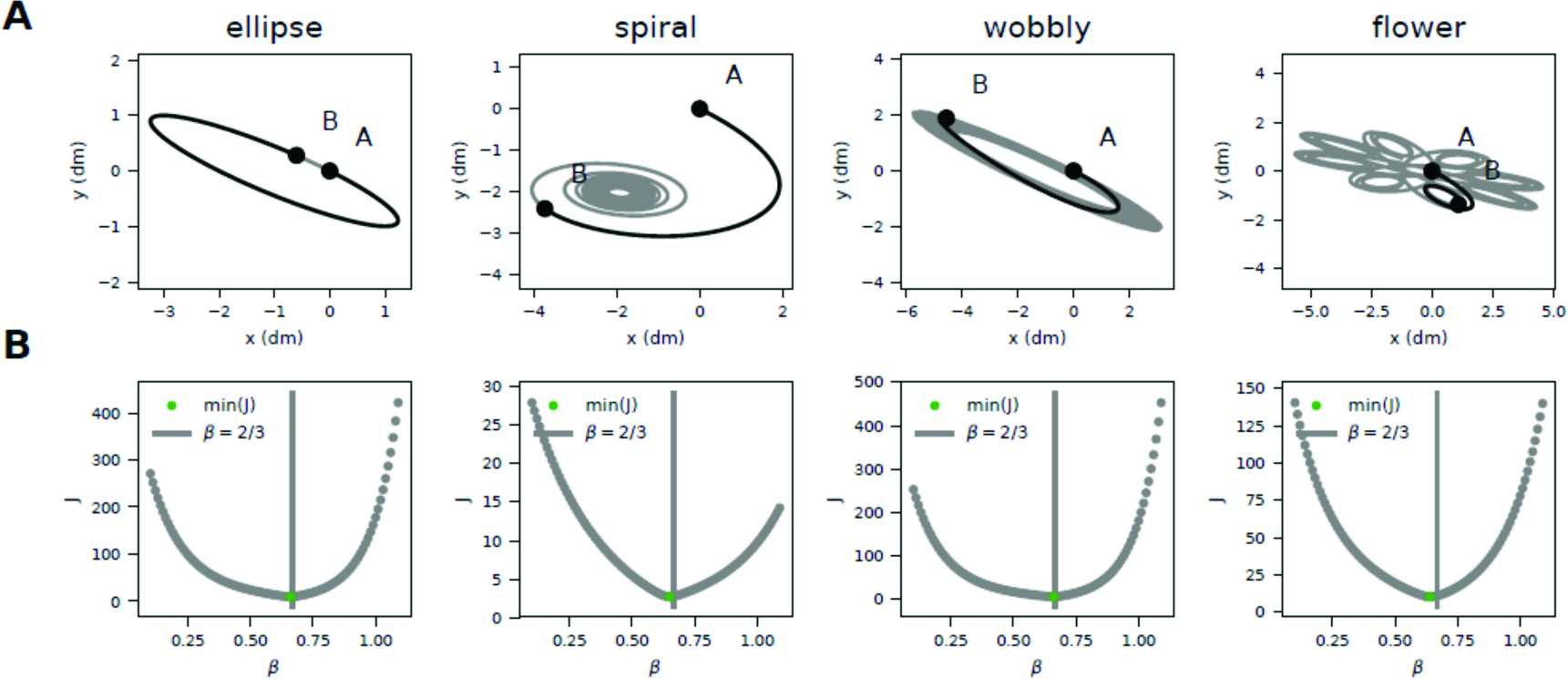
Numerical estimation of minimum jerk for the four quasi-pure (ν=2) frequency curves. (**A**) Trajectory segments analyzed (in black). (**B**) Total jerk as a function of different power law kinematics shows a minimum near β=2/3.

So, to end we show that quasi-pure curves with a peak at ν=2, produce minimum jerk when kinematically traversed at β=2/3 (**Figure 6B**). This confirms and expands the findings in (**Huh and Seknowski, 2015**), at the same time that provides a numerical method to estimate and predict the intricate relationship between geometric purity, kinematic scaling and dynamic optimality for any drawn movement beyond (the ultra-studied) ellipses.

### 3.7. The subtle relationship between curve purity, scaling and optimality

Let us end with a fun and illustrative example to recapitulate. Tracing the contour of Homer Simpson’s face (**Figure 7A**) was drawn on an interactive graphics tablet, tracing the original image shown on the screen, in a single movement, without lifting the pen from the screen. The raw data are smoothed before analysis (see **Methods**). The X and Y coordinates over time (**Figure 7B**) show constant movement with no breaks. Curvature and velocity look fairly correlated (**Figure 7C**), but do not exactly conform to a power law (**Figure 7D**). In fact, the log-log plot seems to indicate multiple segments with different power law exponents, perhaps related to different segments of the drawing. The geometry spectrum analysis shows multiple peaks at low frequencies, and we can see that this is not a pure frequency curve (**Figure 7E**).

**Figure 7.**
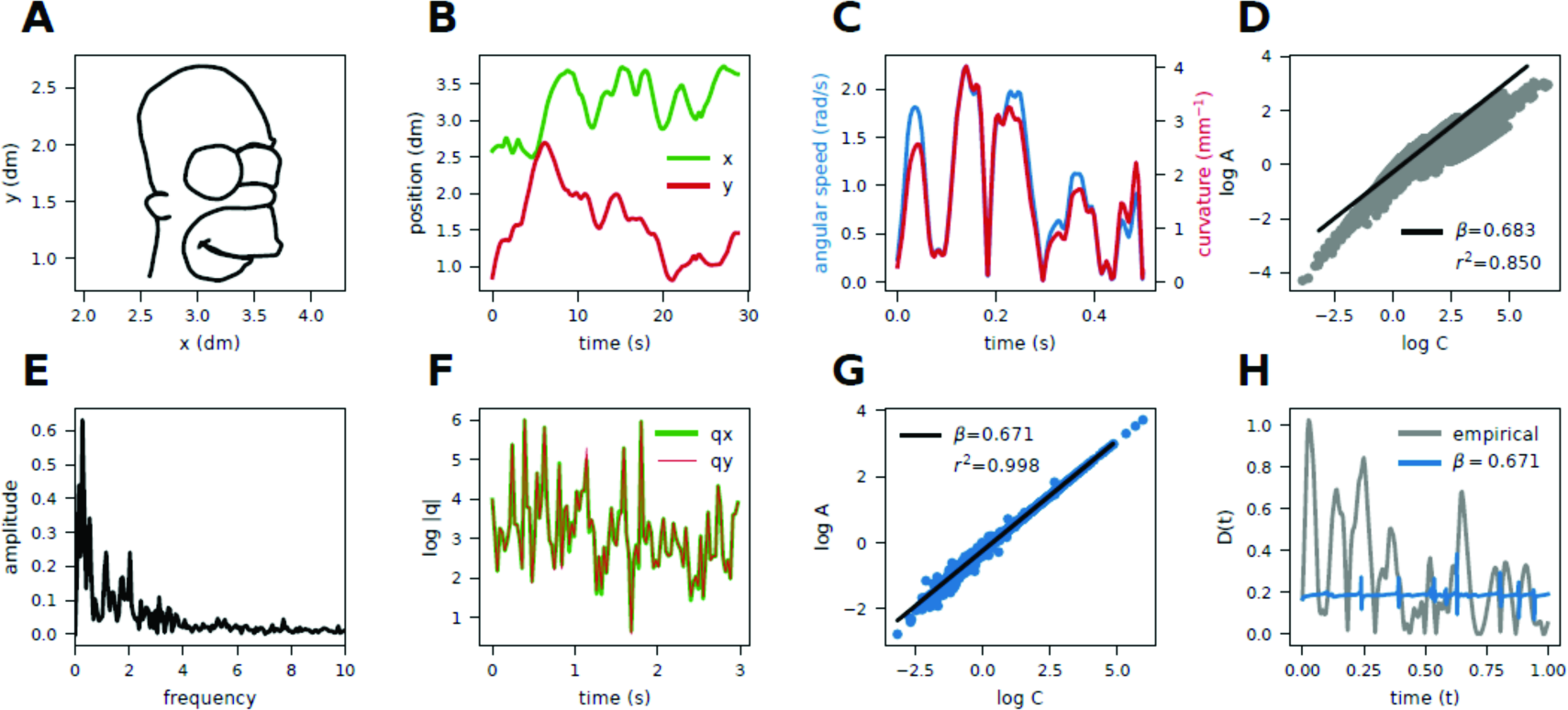
Using Homer’s face to illustrate purity, scaling and optimality in drawing movements. (**A**) Trace of Homer Simpson’s face contours. (**B**) X and Y positions of the tip of the pen as a function of time. (**C**) Speed and curvature for a brief segment a function of time. (**D**) Log-log plot of speed and curvature reveals that kinematics does not comply with a power law. (**E**) Curve spectrum reveals that the drawing is not a pure-frequency geometry, having several peaks at low frequencies and a decreasing tail. (**F**) One can transform the original kinematics so that both X and Y follow the same third-order differential equation with the shared time-dependent parameter q(t), which can be calculated as the ratio between the third and first derivatives of position, shown here. (**G**) Homer’s face now must follow a 2/3 power law. (**H**) The term D, as compared to the original drawing (grey) is quasi constant along the trajectory (blue).

The plots in **Figure 7F-H** show a transformed trajectory: the geometry is the same (still Homer’s face), but the empirical kinematics of drawing are transformed to strictly follow the 2/3 power law (**Figure 7G**). From the same trajectory we can extract the function q(t), and we can see it is near-identical in X and Y dimensions (**Figure 7F**) which, as we saw at the beginning of this article, is a hallmark of a 2/3 power law trajectory. Beyond ellipses, or the other three main curves systematically analyzed in this study, there are infinitely many ways to have a 2/3 power law trajectory (Homer’s face included). As such, the magnitude of the cross product (the term D) is now near constant, unlike the empirical one, which is more variable (**Figure 7H**).

Unfortunately for Homer, since its geometry is not pure (**Figure 7E**), its tracing cannot enjoy both kinematic scaling (**Figure 7G**) and dynamic optimality at the same time. In other words, drawing movements cannot be minimum jerk if speed scales with curvature unless their curvature spectrum is pure. However, in general, one could have minimum jerk using some unknown minimization procedure for any non-pure geometry, with non-power law kinematics.

## 4. DISCUSSION

Nearly forty years later (**Lacquaniti et al., 1983**), the two-thirds speed curvature power law of human movement is still puzzling. Moreover, evidence for the same scaling law with different exponents has recently been discovered empirically (**Huh and Sejnowski, 2015**), and demonstrated to be derivable from normative principles that require the jerk (the time derivative of acceleration) accumulated along the trajectory to be minimal. Along those lines, it had been claimed that the 2/3 speed-curvature power law of movement is a consequence of minimizing mechanical power (**Lebedev et al., 2001**). If so, the power law could be seen as both an outcome of minimum jerk (**Flash and Hogan, 1985**) and “an outcome of least action” (**Lebedev et al., 2001**). That would be interesting if true. However, here we have demonstrated that this is not the case. We have discovered a flaw in the derivation of Lebedev and colleagues, which is due to a basic physics error in interpreting mechanical work. The connection the authors draw between the term D and mechanical work is inexistent. This invalidates the main claim of their paper. Drawing movements complying with the two-thirds power law do not minimize mechanical work.

The origins of the speed-curvature power law remain debated to date. Therefore, we deemed it necessary that the so-far (and to the best of our knowledge) undetected mistake in (**Lebedev et al., 2001**) —and its corresponding unexpected link to equi-affine speed, in the line of the work by Flash and colleagues— does not continue unreported and uncorrected.

However, two pieces of their mathematical treatment are still valuable when expanded upon. They provide more insights to further understand the 2/3 speed-curvature power law observed in humans while drawing. First, their mathematical treatment demonstrates that drawing movements complying with the 2/3 power law must obey a third-order linear ordinary differential equation that only depends on a time-dependent coefficient q(t). The authors explored only the family of x(t) and y(t) solutions when q(t) is constant, which comprises ellipses, hyperbolas and parabolas. Here we exploited some other non-trivial curves of the myriad of geometries that can stem from time dependencies in q(t). Second, the variational principle they put forth demonstrates that D is minimal when it is constant. We tested it numerically, and reformulated it to show that equi-affine displacement is invariant upon different power-law and non-power-law kinematics. We also demonstrated that β=2/3 power laws with ν=2 beyond ellipses have minimum jerk.

Our work has limitations. First, note that except the hand-drawn ellipse and Homer’s face, the rest of our analysis is based on mathematics and numerically simulated curves. Further studies should mirror our findings to experiments inspired by them. Second, all our numerical estimates regarding minimization demonstrate local, but not global, minima. Although it is unlikely, we cannot numerically rule out that a very particular kinematics beats the 2/3 power law when it comes to optimizing the functional of D, V_EA_ or Jerk. Third, a very interesting aspect remains fairly unexplored: while the equation that generates all possible 2/3 power law movement trajectories is a third-order differential equation, in physics virtually all equations of motion do not go beyond second-order. Fourth, while in most human traces and drawings one constantly switches from clockwise to counter-clockwise movement, all curves explored in this manuscript (except Homer’s) where monotonic in curvature. Fifth, it is still a challenge to robustly estimate jerk from empirically measured trajectories because of sensitivity to filtering and to noise in the derivatives.

To end, let us emphasize that the discovery of non-trivial constraints in nature —like a power law— is always as puzzling as rewarding. Kepler established one for the motion of planets. Lacquaniti and colleagues found another one for the movement of hands. Both characterized by an exponent whose value is exactly 2/3. In 1981 Yoshio Koide uncovered a yet-unexplained relation between the masses of three elementary particles (the three charged leptons: the electron, the muon, and the tau): their sum divided by the square of the sums of their square roots is approximately equal to 2/3. If that wasn’t enough, the same relation holds for the masses of the three heaviest quarks. It is tempting to dismiss such phenomenological discoveries as mere numerology or, at best, as simple descriptions awaiting for the hard-core science to take place. This is even more so in biology, where “mechanism” is king while “phenomenon” often enjoys negative connotations. Be it as it may, phenomena borrow from mechanisms that reasons by which they are explained, and restore them to mechanisms in the form of scientific questions which they have stamped with their own meaning. Or, put plainly, the depth that the answer provides very much depends on the quality of the question asked in the first place. Good science is, in a sense, like in good journalism.

## Supplementary material: code and data availability

All code and data used in this study are available. The Jupyter notebook at Google Colab contains numerical simulation code, as well as ALL the analysis and software to reproduce the plots of this manuscript: https://colab.research.google.com/drive/19PkOy4eSsPZIyNqv1o2zXs_BHOCk3HpY. The Github repository contains the copy of the same notebook, the python code for recording of image tracing, the raw data from a participant tracing a Homer drawing, the raw data of ellipse tracing, and video files of animations of changes in D and P in the ellipse and spiral trajectories: https://github.com/adam-matic/purity_scaling_optimality

## Contributions

Idea and conceptualization: AGM; analyses: AM & AGM; mathematical calculations: AGM; figures: AM & AGM; manuscript: AM & AGM.

## Acknowledgements

We thank Regina Zaghi-Lara for drawing Homer’s face.

